# Neural crest mural cells of forebrain meninges harbor innate immune functions during early brain development and exhibit different responses to septic and toxic insults

**DOI:** 10.1101/2023.11.28.569020

**Authors:** Diego Amarante-Silva, Emmanuel Bruet, Rémy Gars, Margaux Piechon, Tatiana Gorojankina, Jérôme Bignon, Sophie E. Creuzet

## Abstract

In vertebrates, the cephalic neural crest (CNC), an embryonic multipotent cell population, gives rise to meningeal derivatives, covering and protecting the developing forebrain. In postnatal life, the meninges exert a prime role in developing and maintaining the central nervous system’s homeostasis and help modulate the immune capacities of resident cells and the blood-brain barrier. However, their role in the embryo remains elusive before the formation of brain barriers. Here, we explore whether meningeal CNC-mural cells participate in immunosurveillance during the early stages of embryogenesis. Using primary cultures of E8 chicken forebrain meningeal mural cells, immuno-phenotyping, and flow cytometry analyses, we demonstrated the expression of specific receptors, including TLRs and MHC class II molecules typically expressed in myeloid cells. Our experiments showed that when challenged by bacterial particles, CNC mural cells also have phagocytic capacities. Transcriptomic analysis revealed activation of inflammatory pathways leading to increased expression of myeloid surface markers and the production of cytokines and interleukins in the presence of bacterial particles. Our in vitro assays also show the capacities of mural cells to present and transfer antigen particles and recruit cooperant cells, both locally and at a distance. Moreover, we revealed that pericytes could deploy DNA traps like neutrophils to stop pathogens. We show that mural cells use extracellular DNA shedding against pathogenic conditions and communicate with each other to mobilize cell cooperation. Transcriptomic analyses revealed that exposure to bacterial particles leads to severe impairment of morphogenetic and neuro-developmental pathways and shed light on the possible detrimental effect of bacterial infection on brain development and maturation of cognitive functions. To reinforce the possible link between neurodevelopmental defects and immune activation, we challenged the immune capacities of CNC pericytes after exposure to toxic medication. We used Valproate (VPA), an antiepileptic drug associated with developmental disorders in newborns when administered during pregnancy. We show that VPA injection into chicken embryos triggered exacerbated myeloid traits and DNA trap releases in CNC pericytes. Our studies show that CNC mural cells are professional innate immune cells capable of pathogen recognition and antigen presentation. This notion breaks the prevailing concept that only mesodermal lineage can participate in the innate immune defense. It opens new avenues for elaborating brain-specific immunity before birth, which may have important biomedical implications for neurodevelopmental disorders.

## Introduction

The neural crest is a phylogenetic innovation which has emerged at the dawn of vertebrate evolution and which turned out to be essential for vertebrate radiation [LeDouarin, 1982; Gans and Northcutt, 1983; Hall, 2000; Green et al., 2015]. During vertebrate development, the neural crest cells give rise to multiple cell types with distinct differentiation potential along the anteroposterior axis. While in the trunk, the neural crest mostly provides Schwann cells and melanocytes, the cephalic neural crest (CNC) gives rise to a rich and versatile ectomesenchyme endowed with skeletal, connective, perivascular, and meningeal differentiation capacities [LeDouarin et al., 2004; Creuzet, 2009a; LeDouarin and Creuzet, 2009]. During chordate evolution, the emergence of CNC-derived mesenchyme has supplanted the cephalic mesoderm in providing dermal, meningeal and skeletal, mesenchymal derivatives to sculpt the vertebrate head [Couly et al., 1992]. Therefore, a developmental shift occurs around the transverse level of the adenohypophysis, between the anterior cephalic domain, where the CNC mesenchymal contribution predominates, and the rest of the body, where mesodermal contribution prevails [Couly et al; 1992; Couly et al., 1993; Creuzet, 2009b]. The concomitant emergence of the CNC-derived ectomesenchyme and the expansion of the prosencephalic vesicles pointed to decisive functional advantages yielded by the CNC cells through the derivatives they form, and presumably also, the mechanisms they control. Developmental studies have shown that CNC cells empower the development of the anterior brain: they provide pro-encephalic signaling to promote encephalization, and convey morphogenetic signaling to coordinate the forebrain patterning [Creuzet, 2009; Garcez et al., 2014; Aguiar et al., 2014].

Anteriorly, the CNC forms the entire muscular-connective tissue wall of the large arteries down to the heart, where they contribute to the semilunar valves, the conotruncus, and the inter-ventricular septum. In the ascendant vascular system, while the endothelium is made of mesodermal cells, the mural cells — pericytes or perivascular cells — have an exclusive neural crest origin [Couly et al., 1995; Etchevers et al., 2001; Korn et al., 2002]. In this ascending vascular sector, emerging from the aortic outflow tract, the CNC-mural cells line the endothelium of the aorta, of the great arteries emerging and its brachio-cephalic branches, to the very thin ramifications of the vasculature which irrigate the face, the oro-nasal, pharyngeal, tracheal mucosae, and the pulmonary veins [Etchevers et al., 2001]. CNC-mural cells also contribute to the vasculature of the anterior brain. In the anterior brain, the CNC is the source of the primary leptomeninges wrapping the forebrain, where CNC cells predominantly differentiate into mural cells that line the endothelium of pial blood vessels and capillaries, and also form the perivascular cells in the forebrain parenchyma — including olfactory bulb, hemispheres, thalamus, epi- and hypothalamus—, along with the stroma of choroid plexus of the lateral and third ventricles [Etchevers et al. 2001; 2002]. They ensure the tuning adjustment of the vascular wall to the hemodynamic constraints and changes in blood pressure imposed by the pulsatile systolic blood supply to support forebrain viability [Etchevers et al., 1999; Creuzet et al., 2002].

At a given transverse level, the mural cells share a common embryonic origin [Etchevers et al., 2001] and similar properties and molecular markers, like the accumulation of alpha-Smooth Muscle Actin (aSMA) [Skalli et al., 1986; Alarcon-Martinez et al., 2018]. Other cytoskeletal proteins such as nestin, tropomyosin, non-muscle myosin, desmin, and vimentin participate in their functions as smooth muscle cells. However, they are not exclusive markers, and their accumulation may vary between organs [Van Dijk et al., 2015]. In addition, Platelet-Derived Growth Factor Receptor (PDGFR-β) is regarded as a specific marker since animals deficient in PDGFR-β show an absence of pericytes, defects in vascularization, and die during development [Lindahl et al., 1997]. However, some authors pointed out that PDGFR-β is widely expressed throughout the embryo and precludes its relevance for developmental studies, but is extensively used to analyze the fate of mural cells in adulthood [Guimaraes-Camboa et al., 2017; Prazeres et al., 2018]. The chondroitin sulfate proteoglycan 4, known as Neuron-Glial Antigen 2 (NG2), also characterizes activated mural cells [Ozerdem et al., 2001]. However, NG2 is widely expressed by other embryonic derivatives like adipocytes, chondroblasts, and Schwann cells, and by oligodendrocytes and microglia, which prevents its extensive use for developmental studies [Fukushi et al., 2003; Castro et al., 2020; Armulik et al., 2011].

As smooth muscle cells, mural cells regulate blood vessel compliance by controlling vessel diameter, from vasoconstriction to vasodilation, and adjusting pro-angiogenic signaling to hypoxic conditions. They exert critical regulatory functions in vasculogenesis by controlling endothelial proliferation, vascular permeability, and vascular matrix assembly [Gaengel et al., 2009; Stratman et al., 2017; Andrews et al., 2018]. Through their mechanical roles and the dynamics of blood flow they act on, mural cells coordinate the homeostasis of the organs they reside in by timely adapting oxygenation, nutrient supply, and draining of metabolic waste [Birbair, 2018]. In the adult brain, mural cells assume pivotal adaptation at the interface of two fluidic compartments, the blood flow and the cerebrospinal fluid [Greitz, 1993; Bergsneider et al., 1998]. They are in close contact with the endothelium, astrocytic end-feet, microglia, and neuronal parenchyma to form the blood-brain-barrier, a barrier crucial to the central nervous system (CNS) homeostasis [Banks, 1999; Rustenhoven et al., 2017; Ganguli and Chavali, 2021]. Through their strategic position between the CNS and the blood [Daneman et al., 2010; Rustenhoven et al., 2017], they control the permeability of the blood-brain-barrier [Daneman et al., 2010; Hill et al., 2015] and regulate the blood flow in the CNS [Carlsson et al., 2023].

Growing pieces of evidence pointed to the implication of mural cells as modulators of the brain-immune interface [Hurtado-Alvarado et al., 2014] upon inflammation, ischemia, oxygen and nutriment shortage, or trauma [Rustenhoven et al., 2017]. Under inflammatory conditions, pericytes can respond to inflammatory stimuli such as Interleukin-1ß (IL-1β) and Tumor Necrosis Factor (TNF-α), Lipopolysaccharides (LPS), and are then able to secrete inflammatory molecules and Damage Associated Molecular Pattern (DAMP) molecules, exhibit transcriptional changes following immunomodulatory stimuli [Pieper et al., 2014; Umehara et al., 2018; Brown et al., 2019], and can engulf pathogen-coated beads [Balabanov et al., 1996]. In the context of cerebral ischemia, oxygen deficit, or glucose deprivation, pericytes repress perivascular markers and prompt the expression of microglia-like markers [Sakuma et al., 2016]. Upon immune-challenged conditions, pericytes can produce inflammatory molecules, including some cytokines, interleukins (ILs), TNF-α, and interferon-γ (IFN-γ) signals, which lead to the recruitment of cells within the neurovascular unit, particularly astrocytes and microglia, into a pro-inflammatory state [Kovac et al., 2011]. However, depending on the context, pericytes may also secrete anti-inflammatory molecules such as IL-33 and fractalkine to prevent microglia activation and exert a neuroprotective role [Rustenhoven et al., 2017; Hattori, 2022]. Furthermore, upon trauma or infection, they concur to leucocyte extravasation to resorb inflammation or limit and defeat pathogen intrusion [Rudziak et al., 2019; Gribl et al., 2018; Julia et al., 2022].

However, the immune properties of pericyte during embryonic development, before the maturation of the immune system and formation of brain barriers, remains an open question. Furthermore, the immune properties of CNC cells still need to be characterized: the overall concept until now specifies that only mesodermal cells can perform immune functions. Here, we explored whether CNC-mural cells can exert immune surveillance at the early stages of brain development. We sought to understand how CNC mural cells respond to pathogenic stimulation and characterize the arsenal of markers and tools they express when challenged by bacterial particles to protect the brain against harmful pathogens. In addition, we aimed to identify the converging mechanisms of toxic and septic shock on the physiology of meningeal CNC mural cells. To do so, we chose to work with chicken embryos where the respective contribution of the CNC and mesoderm lineages to the cephalic vasculature has been accurately mapped [Couly et al., 1995; Etchevers et al., 1999; 2001; Korn et al., 2002], and the sequential development of lymphoid organs and maturation of the professional immune cells extensively documented [Garcia et al., 2021]. Therefore, we focused our study on E8, bearing in mind that at this stage, microglia and astrocytes are not mature, and the brain barriers do not exist as in adults. Moreover, this model, which develops without placenta, allowed us to scrutinize the early intrinsic immune functions of forebrain meninges without any septic variation conveyed by a maternal environment that could impinge on the immune status of the embryo [Guma et al., 2022].

## Results

### CNC-mural cells exhibit a stable perivascular phenotype

To explore the intrinsic immune capacities of CNC-mural cells, we established the optimal conditions for native meningeal cell primary cultures. Meninges peeled from the surface of the telencephalon and diencephalon of chicken embryos at Hamburger-Hamilton stage 32 (HH32, corresponding to E8) were dissociated in a TrypLE express solution, rinsed, and seeded on fibronectin-coated chamber slides, then grown *in vitro* for 72 hours until reaching confluence (Fig. 1A). First, we analyzed the expression of the perivascular phenotype in CNC-mural cell mass culture by testing the accumulation of αSMA, as a marker of smooth muscle cells through immunocytochemistry [Skalli et al., 1986]. Following a rapid adherence to the substratum, the CNC mural cells formed a dense network of multipolar elongated cells strongly positive for αSMA, sharing long interconnected extensions (S1A Fig). Similar phenotypes resulted from either AlexaFluor^TM^ 405-Phalloidin or AlexaFluor^TM^ 633-Phalloidin after fluorescent staining. Quantification revealed that nearly 100% of CNC-mural cells exhibited a perivascular αSMA - positive profile in primary cultures. This phenotype was robust and stable enough to be maintained over ten passages (S1B Fig). In parallel, we tested whether CNC-mural cells also expressed two other mural cell markers, PDGFRß and NG2 [Lindahl et al., 1997; Ozerdem et al., 2001]. Both PDGFRß-(S2A Fig) and NG2-(S2B Fig) positive immunolabeling approached 100% of cells, further confirming the perivascular and pericyte identity of CNC-mural cells.

**Figure 1:**
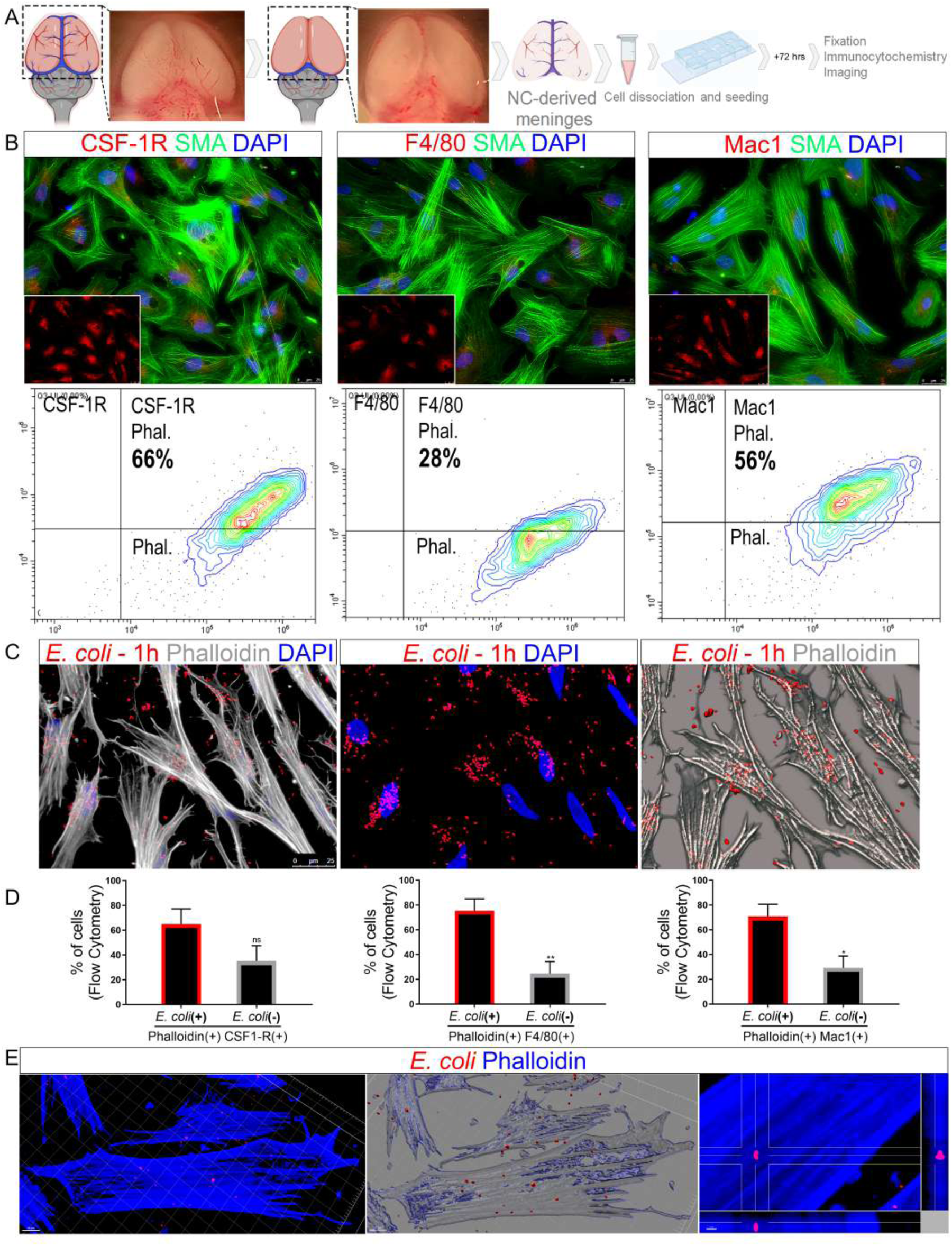
CNC-mural cells harbor immune and phagocytic capacities. (A) Experimental design. Brain from 34HH stage chicken embryo contrasting the prosencephalon with the vasculature and the meninges covering it. The meninges were harvested and dissociated in TrypLE. After cell dissociation, cells were seeded and maintained in culture plates for 72h before fixation, immunostaining, and imaging. (B) Immunostaining showing αSMA-positive (in green) CNC-mural cells expressing macrophage markers (CSF-1R, F4/80, and Mac1respectivily (red) in vitro, in the control condition. DAPI stains nuclei in blue, and insets highlight the expression of macrophage markers alone. The numbers correspond to the mean percentage of double-positive cells quantified with ImageJ software. Below are each marker’s cytometry analysis and the mean percentage of double-marked cells (pericyte-specific marker phalloidin and macrophage markers). (C) CNC-mural cells (Phalloidin-stained; in gray) phagocyte *E. coli* particles (red) in vitro after only 1 h of exposure to the bacterial particles. DAPI stains nuclei in blue. (D) From the left to the right, flow cytometry quantification of Phalloidin-stained cells and positive for the macrophage markers, CSF1-R (64.88%), F4/80 (75.34%) and Mac1 (69.46%), having engulfed fluorescent *E. coli* particles (bar outlined in red), in comparison to *E. coli*-free cells (bars outlined in grey). Error bars represent ± SEM. **p ≤ 0.01; *p ≤ 0.05; ns not significant. (E) Higher magnification of CNC-mural cells showing engulfed *E. coli* red fluorescent particles inside the cell. For a detailed dynamic 3D rendering of a video presented in S04 Fig).

### CNC-mural cells display some surface myeloid markers and harbor phagocytic capacity

To test if CNC-mural cells harbor myeloid features, we examined the expression of early myeloid markers by immunocytochemistry and cytometry, CSFR1 corresponding to CD115, which consists in the earliest marker expressed in macrophage cell lineage [Garceau et al., 2010], F4/80, a macrophage-specific glycoprotein, identified in microglia [Lawson et al., 1990], and Mac1, corresponding to an integrin subunit, CD11b/CD18, identified as a macrophage differentiation antigen [Springer et al., 1979]. Co-immunostaining of primary cultures with these macrophage markers and αSMA antibody revealed that, among αSMA -positive cells, 73% were CSF1R-positive, 44% positive for F4/80, and 47% positive for Mac1 (Fig. 1B). Flow cytometry corroborated these results by showing that, among Phalloidin-stained cells, 66% were CSF1R-positive, 28% F4/80-positive, and 56% Mac1-positive (Fig. 1B). The immunocytochemistry and flow cytometry analyses indicated that the majority of CNC-mural cells exhibited the essential macrophage markers, CSF1R, F4/80, and Mac1, suggesting that they could harbor some immune capacities.

Since the CNC is the exclusive source of the prosencephalon leptomeninges, at odds with the rest of the CNS meninges, we examined whether mesoderm-derived meningeal cells have similar myeloid traits by isolating and culturing meningeal cells taken from E8-spinal cords (S3A,B Fig). We evaluated the expression of macrophage markers by immunocytochemistry and found that the percentage of spinal cord mural cells positive for CSF1R, F4/80, and Mac1 was lower than their CNC-derived counterparts. These observations indicated that spinal mural cells also express macrophage markers, albeit to a lesser extent, and suggested that the CNC-mural cells have a more significant immune potential (S3A,C,D Fig).

Next, we explored the immunological potency of CNC-mural cells by challenging their response to pathogen exposure. To test their ability to phagocytosis, we exposed CNC-mural cells to pHrodo Red *E. coli* BioParticles. This conjugate showed no fluorescent signal at a neutral pH. However, upon internalization, the ingested *E. coli* particles when exposed to the acidic bactericidal pH of the phagolysosome, activated the bright fluorescence of the pHrodo fusion protein.

To investigate if CNC-mural cells harbored intrinsic phagocyte activity, the CNC-mural cells, once reaching the confluence, were exposed to the pHrodo Red *E. coli* BioParticles for 1 h before fixation in 4% formaldehyde, then co-immunostaining with AlexaFluor-633-Phalloidin and DAPI. Confocal imaging revealed that CNC-mural cells had internalized many fluorescent *E. coli* particles in their cytoplasm and filopodia (Fig. 1C). Some fluorescent *E. coli* bacterial particles were also anchored to the cell membrane and accumulated at the cell surface. Interestingly, despite ingesting many *E. coli* particles, the phagocytic CNC-mural cells did not acquire ameboid morphology. Still, they kept their elongated multipolar morphology and formed a dense network with long branching processes. Flow cytometry analysis allowed the quantification of phagocytic capacities in these cells. It showed that most CNC-mural cells expressing macrophage markers, with the respective proportion of 64.88%, 75.34%, 69.46% of CSF1-R-positive, F4/80-positive, or Mac1-positive cells, could phagocyte and internalize *E. coli* particles (Fig. 1D,E; S4 Fig).

### CNC-mural cells have a specific transcriptomic immune signature

To further characterize the immune profile of the CNC-mural cells, we performed a pairwise analysis comparing the transcriptome of *E. coli*-challenged cells to the control ones. We explored changes in gene expression in the presence or absence of treatment with *E. coli* by RNA-sequencing. When challenged by *E. coli* particles, CNC-mural cells showed variations in the expression of more than 600 genes (Fig. 2A). Gene ontology analyses revealed that transcriptomic enrichments coincided with inflammatory responses and cytokine activation (S5 Fig). First, we evidenced the up-regulation of TLR4, a sensor of LPS, and significant enrichment in TLR15, a receptor unique to birds and reptiles [Boyd et al., 2012], which activation accounts for the virulence of pathogen proteases [de Zoete et al., 2011]. Even though only subtle and non-significant changes impacted the expression of MHC class I and class II antigens, many targets belonging to the TNF pathway, involving TNF-interacting proteins, *TNIP2* and *TNIP1*, along with TNF-induced proteins, *TNFAIP3, TNFAIP6* and *TNFAIP2*, were strongly up-regulated. Transcriptomic variations in C-C motif ligands revealed significant enrichment in *CCL20*, *CCL4*, *CCL17*, and *CCL19* chemokines. In addition, *IL1B*, *IL8*, *IL6*, *IL17C*, and *IL18* were significantly augmented in CNC-mural cells when exposed to *E. coli* particles. Similarly, *IFN-γ IRF7*, and *IRF1* expression augmented while *IRF6* diminished. Interestingly, the expression of two cytokine-inducible intracellular molecules, *SOCS3* and *SOCS1*, which act as a suppressor of cytokine signaling, known to temper the inflammatory response of resident immune cells in the adult brain [Baker et al., 2009] and to stimulate macrophage polarization [Wilson, 2014], were both significantly augmented (Fig. 2A).

**Figure 2:**
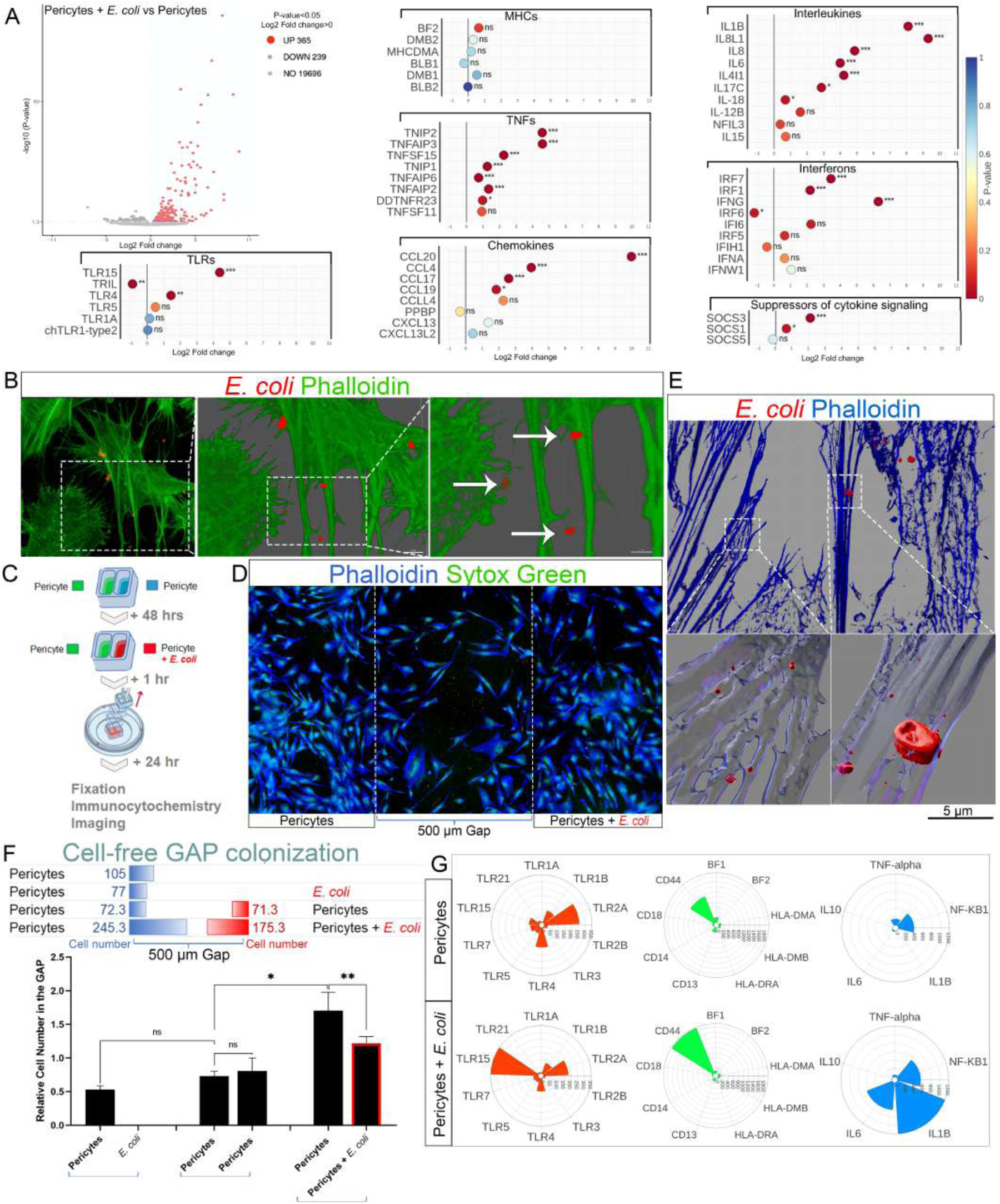
Exposure to *E. coli* particles enhances the immune functions of CNC-mural cells. (A) Volcano plot of Differentially Expressed Genes (DEG) between control and *E. coli* exposed cells, with the upregulated genes shaded in red. Scatter plots displaying the results of RNA-seq analysis of DEG involved in the immune response comparing control and *E. coli*-treated cells. ***p ≤ 0.001; **p ≤ 0.01; *p ≤ 0.05; ns not significant. (A) Immunostaining showing pericytes tagged with the specific marker Phalloidin (green) and *E. coli* particles (red). A 3D reconstruction is shown in the middle panel, and the left panel displays a magnification of filopodial tips with colocalization of *E. coli* particles (indicated by arrows). (B) Representation of the experimental design for the inserts assay. Meningeal pericytes derived from neural crest cells were cultured for 48h in 2-well inserts. *E. coli* particles were added to one well of the inserts, and after 1h, the inserts were removed. The cells were then further cultured for an additional 24 hours before being analyzed, focusing on the 500 µm gap. (C) Immunostaining of pericytes tagged with the specific marker Phalloidin (blue) and the cell nuclei tagged with Sytox^TM^ Green (green). The 500 µm cell-free gap delineated with the dashed lines 24h after insert removal allows quantifying the number of cells that migrated from each insert well (the left side corresponds to control cells, and the right side corresponds to *E. coli*-treated cells). (D) 3D reconstruction of cell interaction originated from the left and right insert-wells within the 500 µm gap, as described in B and C. The lower left panel shows fragmented *E. coli* particles (red) within cells. On the right side, *E. coli*-exposed cells presented fragmented bacteria following pHrodo-*E. coli* phagocytosis. The lower right panel, where phalloidin staining appears in blue, showed both intact and fragmented *E. coli* particles. (E) Quantification of the number of migrating cells in the 500 µm gap. Each line corresponds to a different condition, indicating the mean cell number in the gap. The graph below compares the relative number of cells and shows that untreated pericytes respond to signals produced by *E. coli*-treated mural cells, leading to increased colonization of the gap. This colonization is minus in conditions with *E. coli* alone or *E. coli*-untreated mural cells. Error bars represent ± SEM. **p ≤ 0.01; *p ≤ 0.05; ns not significant. (F) Nanostring analysis of the transcriptional patterns of immune response genes in control pericyte cell culture (above) and *E. coli*-treated mural cell culture (below). Notably, in vitro pericyte cultures can express genes typically attributed to immune cells, reinforcing that these cells possess an immune transcriptional profile. Comparison of expression patterns reveals increased *TLR15, CD44, NF-KB1, IL1B*, and *IL6* in*E. coli*-treated mural cells.

### CNC-mural cells can interact at distance for cell cooperation and antigen presentation

The above results indicated that native CNC-mural cells harbor immune abilities and are capable of pathogens’ recognition and interception, leading to the activation of pathogen-responding pathways. As evoked previously, CNC-mural cells phagocyting pHrodo *E. coli* exhibited many bacterial fragments attached to the outer surface of their cell extensions (Fig. 2B, arrows). To explore how *E. coli*-challenged CNC-mural cells could interact and to investigate if these interactions could underpin antigen presentation, we used a two-well insert system that allowed the cultivation of two pools of cells separated by a silicone barrier (Fig. 2C). Once the cells reached confluence, the silicon barrier removal created a 500 µm-width cell-free gap to become progressively colonized by the converging migration of the cells coming from the two wells (Fig. 2D). CNC-mural cells were seeded either in one well, only, or in both. After 48h of cell expansion, pHrodo *E. coli* particles or fresh vehicle medium were added to the contralateral well. Following the silicone barrier removal, the cells were further grown for 24 hours. After fixation, a combined solution of Alexa Fluor 405 Phalloidin and Sytox^TM^ Green stained at once CNC-mural cell cytoskeleton and nucleus, respectively (Fig 2D). The quantification of the cells in the gap reflected the propensity of the cells to interact at a distance and to migrate when stimulated or not by their environment. First, we tested the ability of CNC-mural cells to migrate into the gap without any cell in the contralateral well. The analysis of the migrating cells in the 500 µm cell-free GAP showed that *E. coli* particles alone were not significantly attractive for the CNC cells (Fig. 2E). Next, we plated CNC-mural cells in both wells and their respective contribution to the colonization of the gap was comparable and limited as previously. However, when *E. coli* particles unilaterally challenged CNC-mural cells before the barrier removal, the number of *E. coli*-treated cells migrating in the gap towards the non-treated ones was 1.4-fold increased over control experiments, suggesting a cooperation between exposed and non-exposed cells. More striking, the non-treated cells massively migrated into the gap, attracted by the CNC-mural cells exposed to *E. coli* particles; their migration and gap colonization was 3.4-fold higher than in control conditions.

These experiments showed that a focal infection stimulated CNC-mural migration and prompted the distant cell interactions between *E. coli*-treated and non-treated cells, presumably underpinning antigen presentation and cell cooperation. To get insight into this process, we used super-resolution imaging and 3D rendering of pathogens phagocyting cells (Fig. 2F). On the exposed side, both entire *E. coli* bacteria and small fragments of bacterial particles co-habited in the cytoplasm and filopodia of phagocyting cells, showing that the CNC-mural cells could degrade pathogens (Fig. 2F; S6 Fig). To confirm the effectiveness and dynamics of *E. coli* bacteria proceeding in the exposed cells, we performed holotomography live imaging with correlative red fluorescence detection of *E. coli* particles in the exposed cells. A 15-minute time-lapse video recording showed the efficiency of the CNC-mural cells in fragmenting the ingested *E. coli* particles (S7 Fig).

On the contralateral side, we did not retrieve any intact bacteria in the cytoplasm of the non-exposed cells. However, many small bacterial fragments were incorporated and internalized to their filopodia, casting light on bacterial material transmission between *E. coli*-exposed and non-exposed cells (S8 Fig). Overall, these observations indicated that the *E. coli* bacteria were degraded and transferred to the non-exposed cells once engulfed by the exposed cells.

To further characterize the effects of *E. coli* exposure on CNC-mural cells and document the molecular determinants underlying the cooperation between the *E. coli*-exposed and non-exposed cells, we performed a refined transcriptomic analysis using a Nanostring-specific code set. This strategy offered a unique, sensitive quantification of the expression of genes of interest based on the amplification-free detection of low-abundance transcripts (Fig. 2G). This analysis revealed that in control native conditions,

CNC-mural cells exhibited a basal expression of *TLR1B, TLR2A, TLR4, TLR5, TLR7, TLR15*, and *TLR21*, the exclusive expression of the MHC Class I, *BF1* factor (no transcript detection for *BF2*), the combined expression of *HLA-DMA* and *HLA-DMB*, a slight expression of *CD13* and *CD18*, and most importantly, *CD44*, a marker shared by CNC and hematopoietic lineages [Corbel et al., 2000; Cao et al., 2016]. When subjected to *E. coli*, CNC-mural cells reduced the expression of TLRs except for TLR15, which was strongly increased. Similarly, while all the CDs and MHC class I and II markers decreased, CD44 expression was 1.5-fold higher than in control cells. Using the same strategy, we explored the expression of cytokines *TNF-alpha, IL1B, IL6, IL10*, and the nuclear factor *NF-KB1*. Under control conditions, CNC-mural cells exhibited a basal expression of *TNF-alpha, IL6*, and *NF-KB1*. However, upon *E. coli* exposure, CNC-mural cells strongly augmented the expression of *IL1B* and *IL6*, along with *NF-KB1* (Fig. 2G). According to the paradigm, when exposed to LPS, CNC-mural cells activated a similar set of immune genes, with a subtle difference in response amplitude to the pathogens or its surface motif. By contrast, exposure to Zymosan demonstrated that CNC mural cells activated a specific immune response the pathogens recognized; the CNC mural cells could activate stereotyped immune responses (S09 Fig). Notably, apart from these inflammatory cytokines, we did not detect any transcript of the counter-regulatory cytokine *IL10* in either *E. coli*-, LPS-, or Zymosan-exposed or non-exposed cells.

### CNC-mural cells can decondense their DNA like neutrophils do

In 2004, Brinkmann et al. reported the capacity of Neutrophils to form DNA traps to intercept and inactivate circulating pathogens [Brinkmann et al., 2004]. Ever since, the capacity to form a DNA web has become an innate immune response against bacterial or fungal intruders was extended to others immune circulating cells. However, this process, so far perceived as the prerogative of adult blood cells, has never been evidenced in mesenchymal cells. Furthermore, the capacity to shed DNA is defective in neonates, casting light on the possible vulnerability of newborn immunity [Yost et al., 2009].

So, to step further towards the demonstration that CNC-mural cells are professional immune cells, we explored if these cells have the capacity to form extracellular traps (ETosis) and shed DNA web.

In the first approach, we performed DNA staining with DAPI of CNC mural cell cultures. Under control conditions, a fraction of the CNC-mural cells, representing about 12%, showed spontaneous decondensation of their nucleus and formation of long of extracellular DNA fragments, released unidirectionally into the extracellular space (Fig. 3A, left panel). When subjected to a septic stress caused by exposure to pHrodo *E. coli* particles, cells exhibited escalating DNA shedding (Fig. 3A, right panel). Under these conditions, the percentage of DNA-shedding cells rose to 25.55% (Fig. 3A, left panel). Furthermore, the quantification of *E. coli* particles bound to extracellular DNA filaments indicated a positive correlation between the percentage of cells exhibiting DNA decondensation and the number of *E. coli* particles in contact with the DNA (Fig. 3A, center graph). In addition, we evidenced that under control conditions, the DNA decondensation was primarily unidirectional, respective to the shedding nucleus. By contrast, upon *E. coli* exposure, the DNA shedding became abounding, tangled and multidirectional (Fig. 3A, left graph). These findings indicated that CNC-mural cells may play a role in the immune response to bacterial infections by generating DNA traps. Bacterial stimuli enhanced the DNA shedding, leading to a distinct configuration of the DNA webs shed in *E. coli*-treated cells and control conditions.

**Figure 3:**
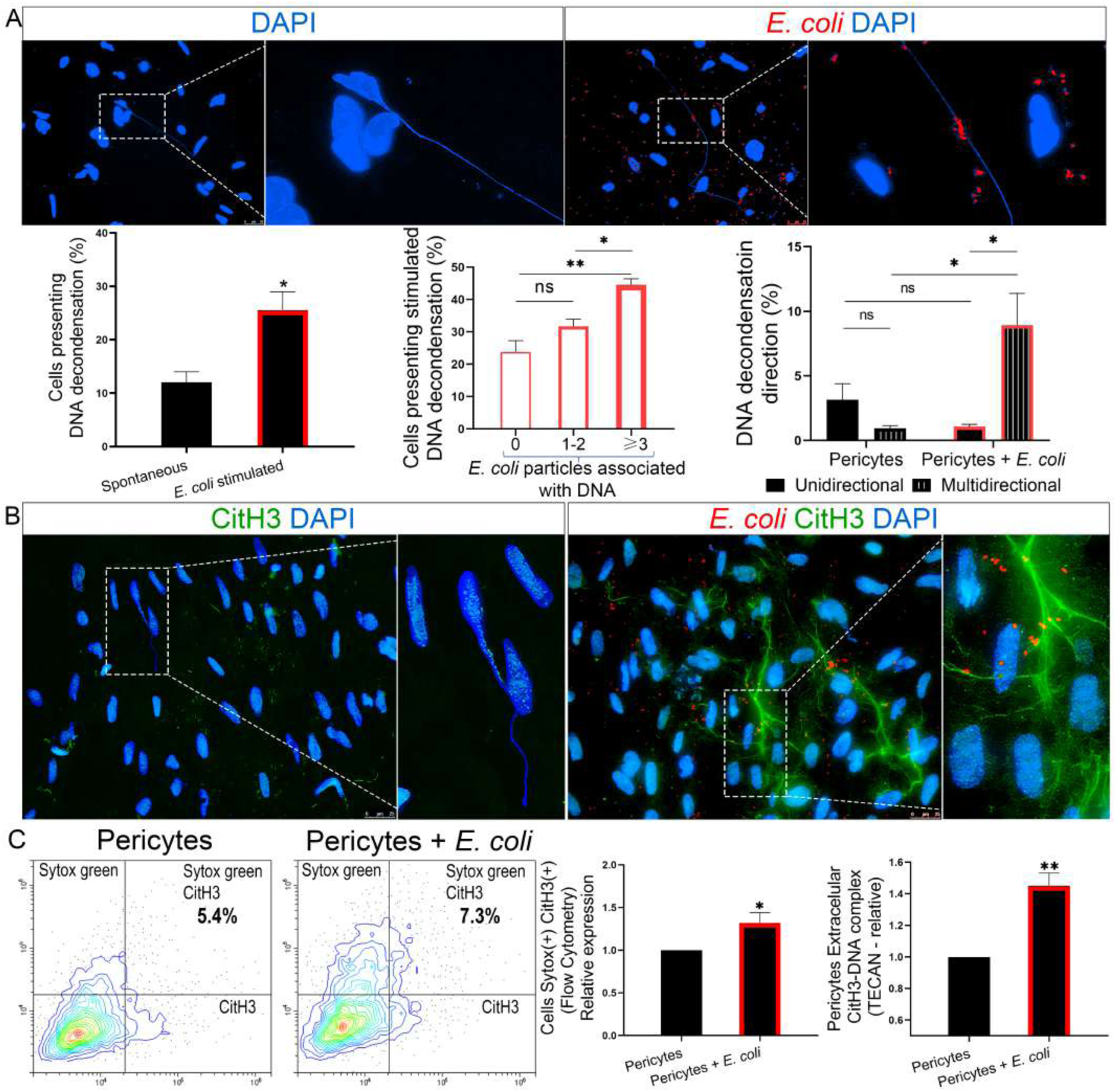
Meningeal NC-derived pericytes present DNA decondensation and produce extracellular traps (ET) in vitro. (A) Pericytes cell culture illustrating DNA decondensation where nuclei appear in blue and phagocyted bacteria in red in both control and *E. coli*-treated (stimulated) cells. The accompanying graphs compare the percentage of cells displaying DNA decondensation in control and *E. coli*-stimulated conditions. The graphs also show the percentage of *E. coli*-stimulated cells displaying DNA decondensation associated with 0, 1-2, or ≥3 *E. coli* particles and the percentage of unidirectional or multidirectional DNA decondensation in control and *E. coli*-stimulated cells. **p ≤ 0.01; *p ≤ 0.05; ns not significant. (B) Mural cells immunostained for CitH3 (green), nuclei (blue), and *E. coli* particles (red) show CitH3 colocalization with decondensed DNA in control conditions. In pericytes treated with *E. coli*, extracellular CitH3 formed a web-like structure associating many *E. coli* particles and other cell nuclei. (C) Double staining of Sytox^TM^ green and CitH3 in unfixed pericytes cell culture treated or not treated with *E. coli* particles. The numbers in the figure represent the mean of double positive cells as determined by cytometry analysis. The first graph illustrates the relative co-expression of Sytox^TM^ green and CitH3 in pericytes treated or not treated with *E. coli*. The second graph represents TECAN analysis results, displaying the relative expression of the CitH3-DNA complex in the conditioned media of pericytes treated or not treated with *E. coli* particles. **p ≤ 0.01; *p ≤ 0.05.

### *E. coli* exposure enhances DNA web shedding from CNC-mural cells

The process of extracellular DNA shedding involves the release of chromatin, initiated by the enzyme Peptidyl Arginine Deiminase (PADI) 4, which induces the citrullination of histones [Thiam et al., 2020]. By reducing their surface charges, citrullinated histones (CitH) fail to maintain cohesive interactions with the chromatin and prompt nucleosome disintegration [Leshner et al., 2009]. We tested the accumulation of CitH3 (bearing three sites of citrullination along their tail on R2, R8, and R17) as an immunomarker of intense citrullination and effective DNA shedding. In our experiments, the accumulation of CitH3 in control cells appeared as faint staining of the nuclei, shedding unidirectional DNA strands (Fig. 3B, left panel). When subjected to *E. Coli* particles, CitH3 immunostaining revealed dense and multidirectional extracellular webs trapping *E. coli* particles and interconnecting many adjacent nuclei (Fig. 3B, right panel). We used Sytox^TM^ Green staining and CitH3 immunodetection to quantify the DNA shedding by cytometry. Due to its incapacity to enter intact cell membranes, Sytox^TM^ Green, a DNA binding dye used for cell viability assays, was a resolutive strategy to evidence the shedding nuclei and extracellular DNA exclusively. We observed that 5.4% of living cells were spontaneously releasing DNA traps, which was significantly 1.3-fold higher in CNC-mural cell cultures treated with *E. coli* particles (Fig. 3C, right panel). In parallel, the quantification of cell culture grown on the Tecan microplate showed that *E. coli* exposure significantly augmented the percentage of adherent cells undergoing ETosis. In addition, when the supernatant of the cell culture treated with *E. coli* particles was harvested and analyzed on the Tecan microplate previously coated with CitH3 antibody, this approach revealed an increase of the complex associating CitH3 and extracellular DNA shed in their supernatant, even higher than at the intra-cellular level (Fig. 3C, left panel).

Overall, our observations support with the demonstration that CNC-mural cells harbor native immune functions early in brain development, suggesting that these capacities may operate as a first line of defense against pathogens at a stage when brain barriers and the cognate professional immune cells are not mature.

### Valproate, a toxic medication for brain development impedes the native immune functions of CNC-mural cells

To explore if this function could be targeted by toxic medications leading to severe post-natal behavior impairment, we challenged the CNC-mural cells with Valproate (VPA), an anticonvulsant drug widely administered to manage seizures, as well as an efficient medication and mood stabilizer alleviating with a broad spectrum of conditions including bipolar disorders and schizophrenia [Löscher, 2002; Bowden, 2009; Emrich et al., 1980]. However, aside from its powerful effects on adult brain disorders, this drug was embryotoxic when administrated during pregnancy. Exposure to VPA during development had severe drawbacks on fetal development, leading to an increased risk of congenital malformations like spina bifida and exencephaly, as well as post-natal behavioral deficits and autistic manifestations [Nau et al., 1991; Christensen et al., 2013].

To investigate the consequences of toxic medication on CNC-mural cells, we treated meningeal cells in the embryo before being collected for in vitro cell culture. In mass culture, the cells showed alterations in the expression of macrophage markers CSFR1, F4/8, and Mac1. For all these markers, we noted an increase of their expressions since 86% of the cells expressed CSFR1, 64% F4/80, and 76% Mac1. Flow cytometry analyses corroborated these results and showed a significant increase for each marker considered. (Fig. 4A) The RNA sequence analysis revealed a transcriptomic profile distinct from the one revealed after pathogenic exposure. Some targets appear significantly increased, such as *TRIL*, MHC class I, and *BF2* (while it is not detected on native cells or exposed to *E. coli*). Similarly, the activation of molecules related to TNF signaling occurred, characterized by an increase in *LITAF, C1QTNF5*, and *TNIP1*, and a decrease in *TRAF3* and *TRAF6*. Exposure to VPA also induced an increase in chemokines, *CCL4*, and *CXCR1*, and a modest increase in *IL6* and some interleukins-related secondary targets (but no activation of *IL1B* and *IL8*, as with exposure to *E. coli* particles). Also, the activation of interferon molecules selectively mobilized *IFI6*, *IFI35*, and *IRF7*. When VPA-treated cells were grown in two-well inserts, we found that they had a restricted ability to migrate spontaneously, either with control cells or with homologous cells also treated with VPA (Fig. 4C). These observations show that VPA prevents the capabilities of CNC-mural cells from interacting and cooperating remotely. On the other hand, the immunostaining performed with Sytox^TM^ Green and CitH3 showed that the treated cells have a significantly higher propensity to deploy their DNA than the control cells. (Fig. 4D). Transcript analysis performed using specific Nanostring code set approach showed that VPA significantly reduces TLR expression on the surface of CNC-mural cells and TNF-alpha signaling. On the other hand, VPA exacerbates CD44 expression while reducing the expression of the self-recognition marker BF1 (Fig. 4E).

**Figure 4:**
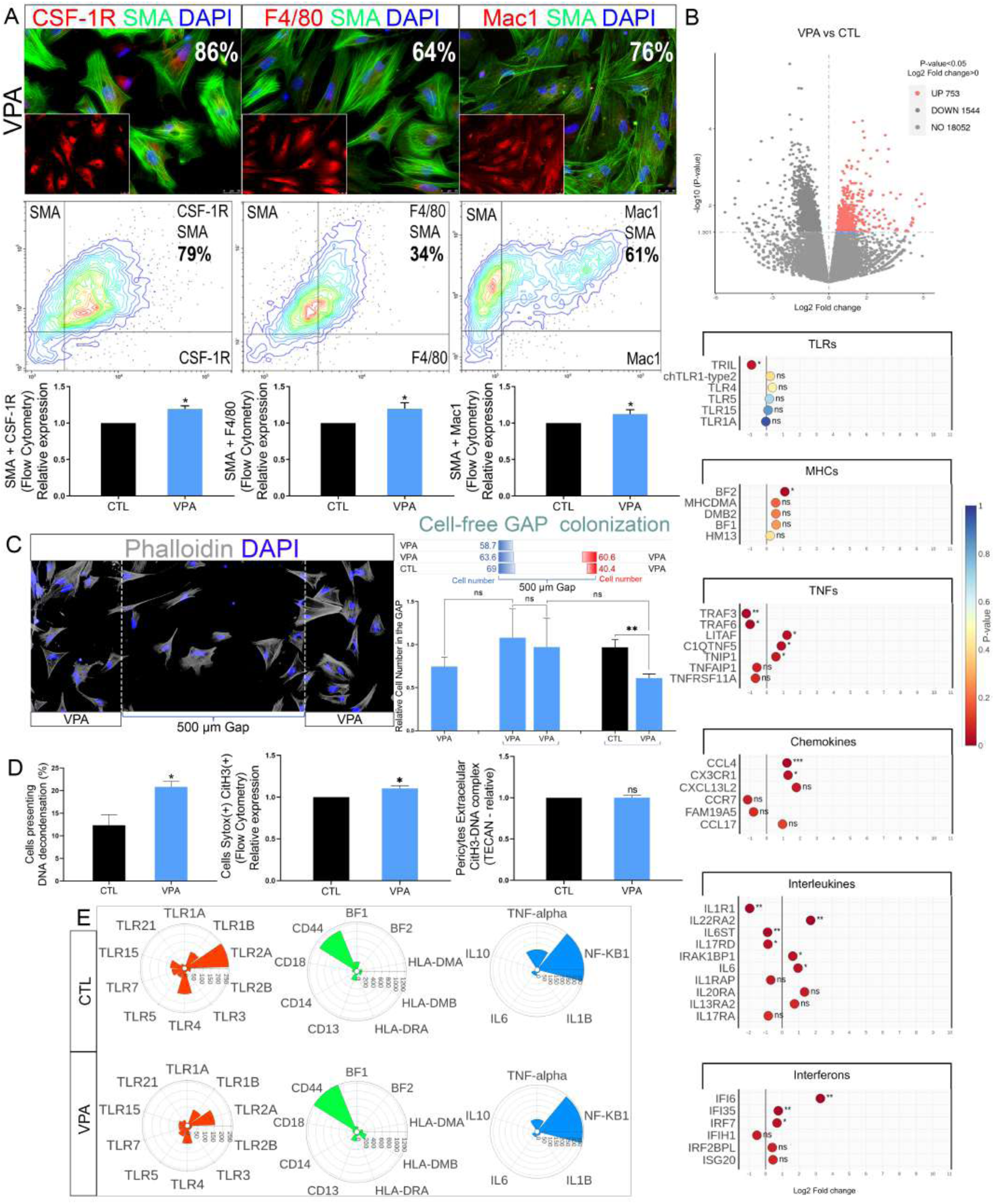
Exposure to VPA modulates the immune profile of meningeal CNC mural cell culture in vitro. (A) Immunostaining showing a αSMA (+) (green) cells expressing macrophage markers (CSF-1R, F4/80, and Mac1 respectively - red) in vitro. DAPI stains nuclei in blue, and insets highlight the expression of macrophage markers alone. The numbers correspond to the mean percentage of double-positive cells quantified with ImageJ software. The center panels below are each marker’s cytometry analysis and the mean percentage of double-marked cells (pericyte-specific marker a αSMA and macrophage markers). The graph panels below compare the relative co-expression of the pericytes-specific marker αSMA and the macrophage markers between control and VPA-treated cells (cytometry). *p ≤ 0.05. (B) Volcano plot of DEG between control cells and VPA-treated cells, with the upregulated genes in red. Scatter plots below display the results of RNA-seq analysis of DEG involved in the immune response comparing control cells and VPA cells*. ***p ≤ 0.001; **p ≤ 0.01; *p ≤ 0.05; ns not significant. (C) Immunostaining of pericytes tagged with the specific marker Phalloidin (gray) and the cell nuclei labeled with DAPI (blue). The 500 µm, large cell-free gap delineated between the dashed lines; the cells migrated during the 24h following dragging the insert. The upper panel on the right quantifies the number of migrating cells in the 500 µm GAP. Each graph line corresponds to a different condition, and the numbers indicate the mean cell number. The graph below compares the relative number of cells found in the 500 µm gap and demonstrates that VPA-treated cells migrate less towards control cells than other VPA-treated cells. Error bars indicate ± SEM. **p ≤ 0.01; ns not significant. (D) The graph on the left compares the percentage of cells displaying DNA decondensation between control and VPA-treated cells. The next one illustrates the relative co-expression of Sytox^TM^ green and CitH3 in control and VPA-treated cells (cytometry). The last graph represents TECAN analysis results, displaying the relative expression of the CitH3-DNA complex in the conditioned media of control and VPA cells. *p ≤ 0.05; ns not significant. (E) Nanostring analysis of the transcriptional patterns of immune response genes in control and VPA-treated mural cells (below) cell culture.

### Dual septic and toxic insults exacerbate inflammatory pathways in CNC-mural cells

When VPA-treated CNC-mural cells were exposed to *E. coli* particles, they demonstrated the ability to phagocyte and present bacterial particles (Fig. 5A). Furthermore, compared to CNC-mural cells from control embryos, the VPA treatment significantly increased the percentage of CNC-mural cells phagocytosing the *E. coli* particles (Fig. 5A).

**Figure 5:**
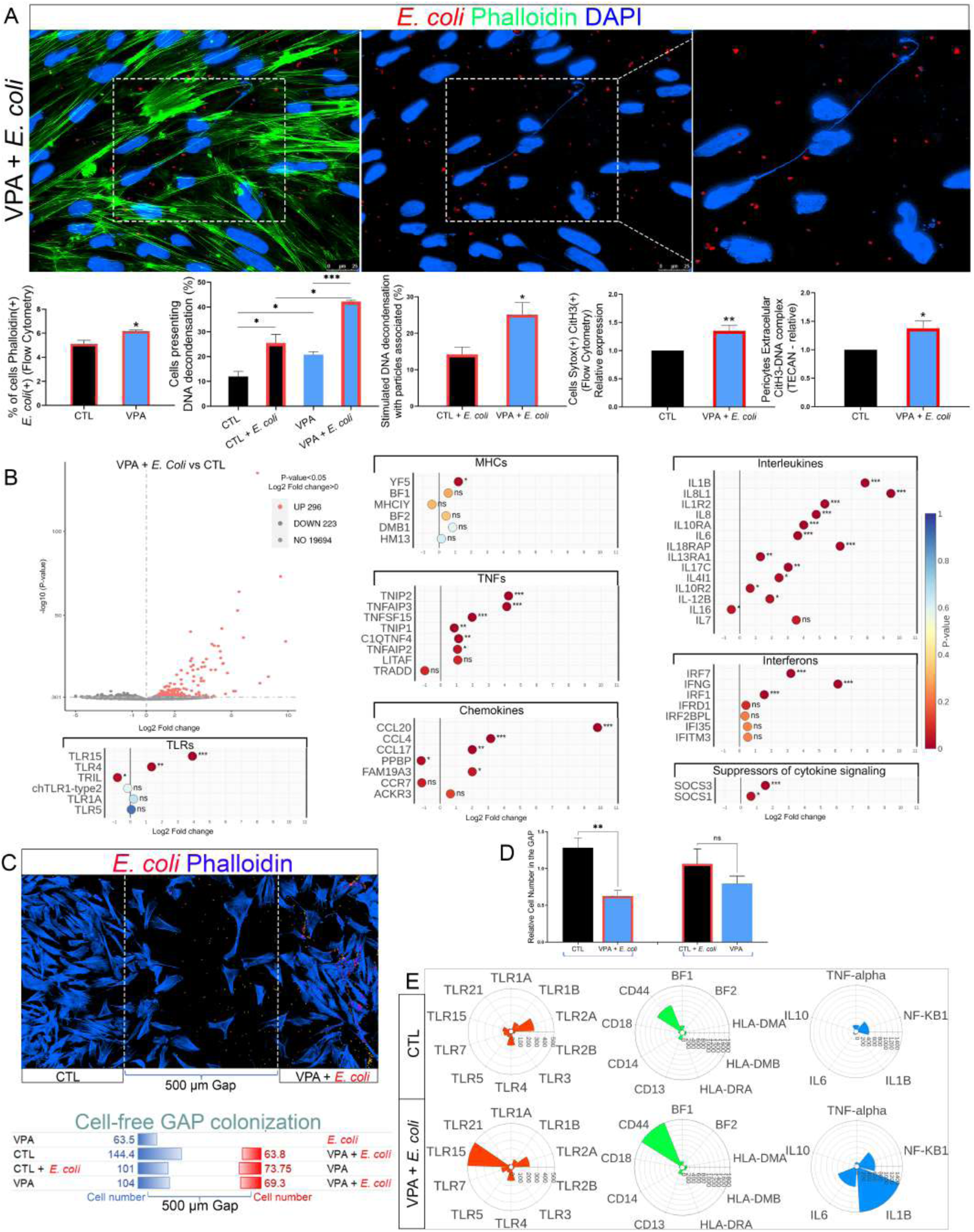
Exposure to VPA and to *E. coli* increases the inflammatory profile of meningeal CNC mural cell culture in vitro. (A) VPA-treated mural cell culture exhibit DNA decondensation after *E. coli* exposure, where cell nuclei appeared in blue and *E. coli* particles were in red. From left to right, the graphs show cytometry quantification of the percentage of control and VPA-treated cells expressing the specific pericyte marker Phalloidin and fluorescent *E. coli* particles after 1h of exposure to *E. coli* particles. The following graph compares the percentage of cells displaying DNA decondensation in control and VPA-treated cells with *E. coli* particles. Then, the graph illustrates the relative co-expression of Sytox^TM^ green and CitH3 in control and VPA- and *E. coli* -treated cells (cytometry). The last graph represents TECAN analysis results, displaying the relative expression of the CitH3-DNA complex in the conditioned media of control and VPA- and *E. coli* -treated cells. ***p ≤ 0.001, **p ≤ 0.01, *p ≤ 0.05; ns not significant. (B) Volcano plot of DEG between control cells and VPA cells* exposed to *E. coli*, with the upregulated genes in red. Scatter plots display the results of RNA-seq analysis of DEG involved in the immune response comparing control and VPA- and *E. coli* -treated cells. The plots show that nearly all genes appeared upregulated after exposure to VPA and to *E. coli* particles. ***p ≤ 0.001; **p ≤ 0.01; *p ≤ 0.05; ns not significant. (C) Immunostaining of control and VPA-treated mural cells tagged with the specific marker Phalloidin (blue) and *E. coli* particles (red). In the gap, the quantification of cells migrated from each insert well; the left side represents the control cells, and the right side represents VPA- and *E. coli*-treated cells. Each graph line corresponds to a different condition, and the numbers indicate the mean cell number. (D) The graph compares the relative number of migrating cells in the gap. It shows that untreated pericytes respond to signals produced by VPA- and *E. coli*-treated pericytes, leading to increased colonization of the gap. Error bars represent ± SEM. **p ≤ 0.01; ns not significant. Nanostring analysis of the transcriptional patterns of immune response genes in control and *E. coli*-treated VPA-treated mural cells. Comparison of expression patterns reveals increased *TLR15, CD44, NF-KB1, IL1B*, and *IL6* in *E. coli*- and VPA-treated cells. (H) Immunostaining of control and VPA-treated. cells with the specific marker Phalloidin (blue) and *E. coli* particles (red). The cells were cultured as described in Figure 2C. The panel below shows the quantification of the number of migrating cells in the gap. Each graph line corresponds to a different condition, and the numbers indicate the mean cell number. (I) The graph compares the relative number of migrating cells in the gap. It shows that untreated mural cells respond to signals produced by VPA- and *E. coli*-treated cells, leading to increased migration to the GAP. Error bars represent ± SEM. **p ≤ 0.01; ns not significant. (J) Nanostring analysis of the transcriptional patterns of immune response genes in control and VPA- and *E. coli*-treated cells. Comparison of expression patterns reveals increased *TLR15, CD44, NF-KB1, IL1B*, and *IL6* in *E. coli*-treated VPA-treated mural cells.

When primed with VPA and exposed to *E. coli* particles, CNC-mural cells exhibited intense DNA decondensation (Fig. 5A). The combination of VPA treatment and *E. coli* exposure considerably enhanced the DNA decondensation and shedding, as 42% of the CNC-mural cells underwent ETosis compared with the control.

In addition to the observed increase in the percentage of pericytes presenting DNA decondensation in response to *E. coli* particles, further analysis revealed that the percentage of cells with *E. coli* particles associated with DNA decondensation was 1.7-fold higher in CNC-mural cells from VPA-treated embryos compared to those from control embryos (Fig. 5A). Flow cytometry analysis showed an increased percentage of VPA-treated cells and subjected to *E. coli* particles, undergoing ETosis, compared to controls (Fig. 5A). Consistent with these findings, the conditioned media of CNC-mural cells from VPA-treated embryos contained significantly more CitH3-DNA complexes (Fig. 5A). These results indicated that CNC-mural cells from VPA-treated embryos exhibit increased DNA decondensation and extracellular DNA web release when exposed to *E. coli* particles.

The RNA sequencing revealed that the CNC-mural cells from VPA-treated embryos had disrupted gene expression in response to *E. coli* treatment. The volcano plot in Fig. 5B highlighted a significant number of up-regulated genes. The analysis of immune response genes showed a similar response to *E. coli*-treated cells from both control embryos (Fig. 2A) and VPA-treated embryos (Fig. 5B).

However, the results of cell migration and colonization of the 500 µm-gap demonstrated that the VPA-treated -CNC mural cells had reduced capacity to migrate and interact with their neighboring cells. These observations suggested that VPA rendered the CNC-meningeal cells less responsive to the inflammatory signals produced by *E. coli*-treated cells (Figs. 5C,D). Altogether, these results suggest that VPA-treated CNC-mural cells can respond to infections but are less attracted to infection or inflammation sites to cooperate.

The Nanostring transcriptomic analysis validated that CNC-mural cells from VPA-treated embryos expressed genes typically involved in the immune response. Furthermore, when exposed to *E. coli*, particles, CNC-mural cells up-regulated several genes, such as *TLR15, CD44, NF-KB1, IL1B*, and *IL6*. These observations indicated that the VPA-treated CNC-mural cells could develop a specific immune response and that exposure to *E. coli* enhanced this response [Fig. 5E].

## Material and methods

### Ethics statement

The Centre National de la Recherche Scientifique (CNRS) and Paris-Saclay University adhere to the ethical use of laboratory animals, under which prehatching avian embryos are not considered live vertebrate animals. As such, this study did not require any animal approval.

### Chicken embryos

The experiments were conducted in chicken embryos. Fertilized eggs acquired from Les Bruyeres (Dangers, France) were incubated at 38±0.5°C with increasing humidity and ventilation until reaching the stages (HH) [Hamburger and Hamilton, 1951] required for experiments.

To test if and how a toxic insult could affect the immune capacities of CNC-mural cells, embryos were injected with a VPA (24 ug /40 mg) or a PBS-vehicle (control) solution into the forebrain region using ultrathin glass needles. After injection and sealing eggshells with tape, embryos were preincubated until reaching the desired HH34 stage to harvest their meninges.

### CNC-mural cell isolation and in vitro culture

In vitro experiments were performed on primary cultures of pericytes. VPA and control-injected chicken embryos heads were collected at HH34 stage and placed in PBS solution supplemented with 0.5 % PS and 0.5% Fungizone. The skin was removed and the meninges from the prosencephalon region were collected into 1.5 ml Eppendorf tubes with PBS (1 embryo/tube). The CNC meninges were dissociated with TrypLE (Gibco) for 30 min at 37°C and pelleted by speed decantation. After counting cells to adjust their concentration, CNC-mural cells were seeded on 4-well chamber Polymer slides (Nunc™ Lab-Tek™) coated with fibronectin (5µg/cm^2^, Sigma-Aldrich), and seeded at the concentration of 15.000 cells /cm^2^. Cells were cultured in DMEM/F-12 medium, containing 10 % FBS, 1 % PS, and 1% Fungizone at 37 °C in a humidified atmosphere with 5% CO2. After 3 days of incubation, 25 μL of pHrodo Red *E. coli* BioParticles (Thermo Fisher Scientific) were added into several wells for 1 hour. Cells were then washed with PBS and fixed with 4% formaldehyde (20 min) for further immunostaining.

### Cell-cell interaction assays

To analyze cell-cell interactions and their cooperation at distance, cell cultures were performed as described above, with some adaptations. After isolation, the CNC-mural cells were seeded in the 4-well plates containing self-inserted culture-inserts, with 2 wells separated by a 500 µm removable silicon wall (Ibidi). During the second day of culture, the *E. coli* particles were added to the cells unilaterally, leaving the cells in contralateral well unexposed. After 1h of treatment, the inserts were removed, the cells were rinsed, and a fresh medium was added. Migration of cells into 500µm cell-free gap was analyzed after further incubation for 24h. Cells were then fixed and stained with Phalloidin blue and Sytox^TM^ Green. The cell migration and interactions were observed in fluorescent microscope and the images were analyzed with the ImageJ software.

### Immunocytochemistry

Briefly, after fixation, cells were washed 3 times with PBS and permeabilized with 0.1% Triton X-100 (Sigma, 15 min). Cells were incubated with primary and secondary antibodies overnight at 4°C. Primary antibodies used include αSMA (MA5-11547; Invitrogen), NG2 (sc33666; SantaCruz), PDGFRß (MA5-15143; Invitrogen), CSFR1 (sc692; SantaCruz), F4/80 (sc-377009; SantaCruz), Mac1 (NB110-89-474; Novis Biotechne), CitH3 (ab5103; Abcam), CitH4 (07-596; Sigma Aldrich). Primary antibodies were detected using anti -rabbit- and anti-mouse-Alexa Fluor conjugated secondary antibodies (Invitrogen). Cell nuclei were stained with 4′, 6-diamidino-2-phenylindole dihydrochloride (DAPI, Sigma) and/or with Sytox^TM^ Green Nucleic Acid Stain (1:30.000, Invitrogen). Cells were imaged with a fluorescence stereomicroscope (Leica DM2000) or a confocal microscope (Leica TCS SP8). The images were quantified using the ImageJ software (National Institutes of Health) and processed with Photoshop software (Adobe).

### Cytometry

Primary pericyte cultures were prepared as described above. After 3 days in vitro, the cells were detached from the culture plates with TrypLE (Gibco), counted and approximately 1 x 10^5^ cells were transferred into 1.5-ml tubes. The tubes were maintained in ice during all the procedures before the FACS analysis. Cells were incubated first with primary antibodies overnight at 4°C and then with secondary antibodies overnight at 4°C. For identification of living cells co-expressing Cit-H3 and Sytox^TM^ Green, cells were incubated with agitation with primary anti-Cit-H3 antibody (1:250) for 1 h, washed then further incubated for 1 h with secondary antibodies (goat anti-rabbit IgG Alexa Fluor 633, 1:500). Finally, the Sytox^TM^ Green (1:30.000), a plasma membrane-impermeable DNA-binding dye, was added for 15 min and cells were subjected to FACS analysis. The cells were analyzed with CytoFlex (Beckman-Coulter) bench Flow Cytometer and the data were processed using the CytExpert software.

### DNA and Cit-H3-DNA complex quantification

Primary pericyte cultures were performed as described above, with some modifications to analyze the culture’s supernatant. On the second day of culturing, the medium was changed and cell incubation continued. On the third day the cell cultures were treated with 25µl *E. coli* BioParticles (pHrodo, Thermo Fisher Scientific) alone or combined with a PAD inhibitor Cl-amidine (200µM, Calbiochem, Merk) for 1 hour [Chaaban, et al., 2020]. This cell-permeable compound acts as an inhibitor of DNA decondensation and extracellular DNA trap formation. Then supernatants were collected into the Eppendorf tubes and centrifugated to eliminate cell debris (300XG for 5 min). The DNA concentrations in the supernatants was determined by NanoVue nanodrop (GE Healthcare). Supernatants were stocked at -20°C before further utilization. The cells recovered from these experiments were used for the above-described living cells FACS analysis.

DNA traps are accompanied by CitH3 and CitH4. To detect the DNA quantity associated with CitH3 in collected supernatants, the latter were incubated overnight at 4° in 96 Well Black Plate (Ibidi) previously - incubated overnight with anti-Cit-H3 antibody. The day after, the wells were treated with the nucleic acid–binding fluorescent reagent Sytox^TM^ Green to access the Cit-H3-DNA complexes. The plates were then read using a Tecan Genios Microplate reader, with excitation at 485 nm and emission measured at 535 nm. Results are expressed in arbitrary fluorescent units. Supernatant samples treated with DNAse (20U/mL) served as negative control.

### Transcriptomic analysis with specific Nanostring code-set and bulk RNA-seq

The pericyte cultures were performed as described above, and the total RNA from control, VPA and *E. coli*-treated samples was extracted using the Tri Reagent (Sigma-Aldrich) according to the manufacture instructions. The samples were sent to the genomics platform of the Curie Institute for Nanostring nCounter analysis. The panel of analyzed genes is shown in the supplemental table. Analysis of differential gene expression was performed using the nSolver 4.0 software.

For the RNA-Seq analysis, the RNA samples were extracted in a similar way and were sent to the Novogene Co., LTD (Cambridge, England). Genes with adjusted p-value ≤ 0.05 and log2 (FoldChange)

> 0 were considered as differentially expressed.

### Statistical Analysis

Statistical differences between samples were evaluated by unpaired Student’s t-test using GraphPad Prism 8 software (GraphPad Software, Inc.). The level of significance was set at p ≤ 0.05. The results represent the mean of at least three independent experiments.

## Abbreviations

aSMA: alpha-Smooth Actin Muscle
CitH: Citrullinated Histone
CNC: cephalic neural crest
CNS: central nervous system
DAMP: Damage Associated Molecular Pattern
E: embryonic day
ET: Extracellular Traps
HH: Hamburger-Hamilton stage
IFN: interferon
IL: interleukin
LPS: Lipopolysaccharides
PADI4: Peptidyl Arginine Deiminase 4
PDGFRβ: Platelet-Derived Growth Factor Receptor β
NG2: Neuron-Glial Antigen 2
R: arginine convertible into citrulline, Tumor Necrosis Factor
VPA: Valproate.

