## Supplementary figures and images for "Neural crest mural cells of forebrain meninges harbor innate immune functions during early brain development and exhibit different responses to septic and toxic insults"

### Supplementary Fig1

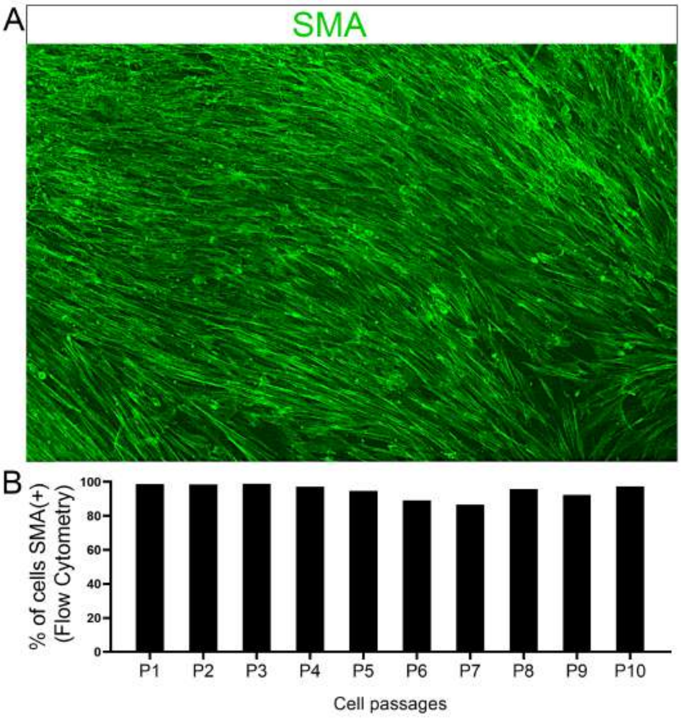

### Supplementary Fig2

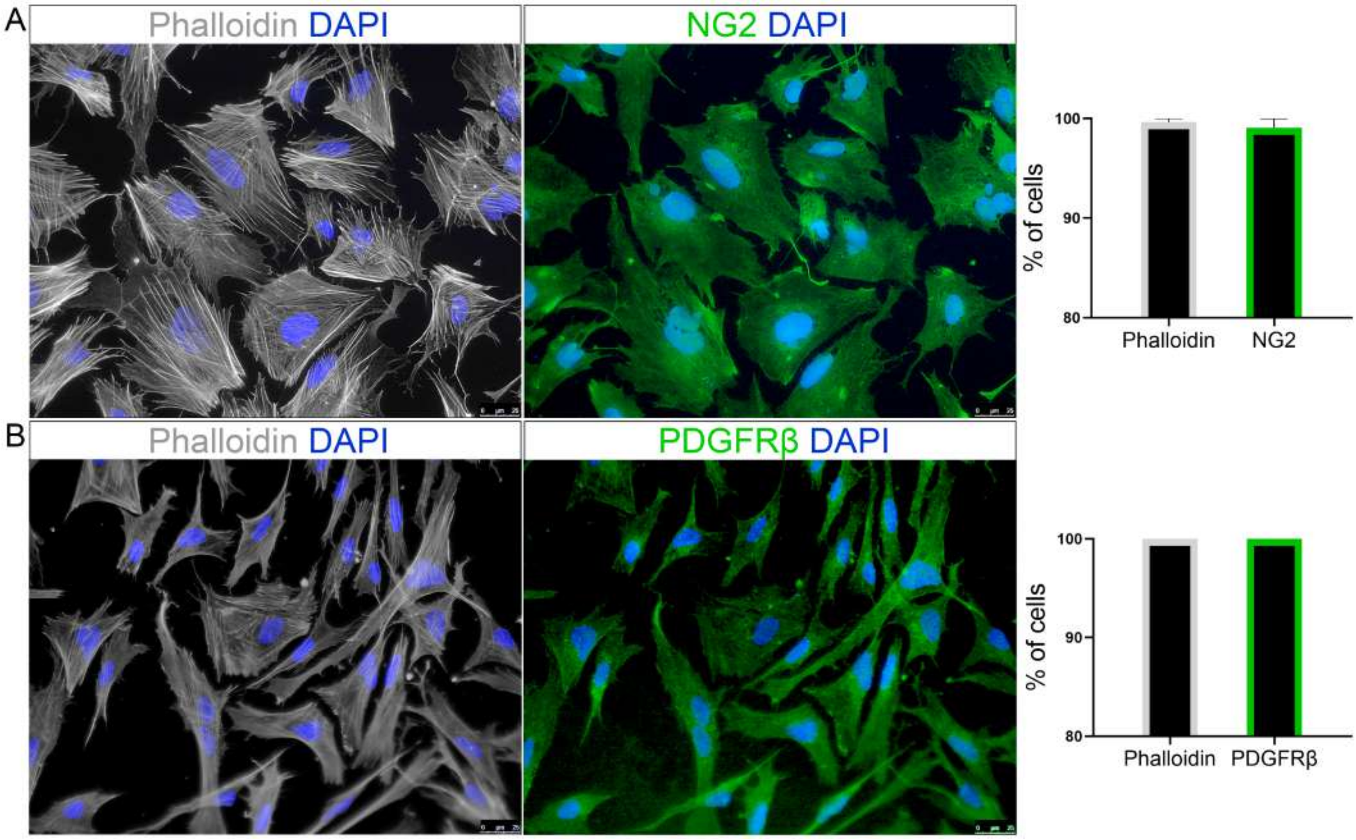

### Supplementary Fig3

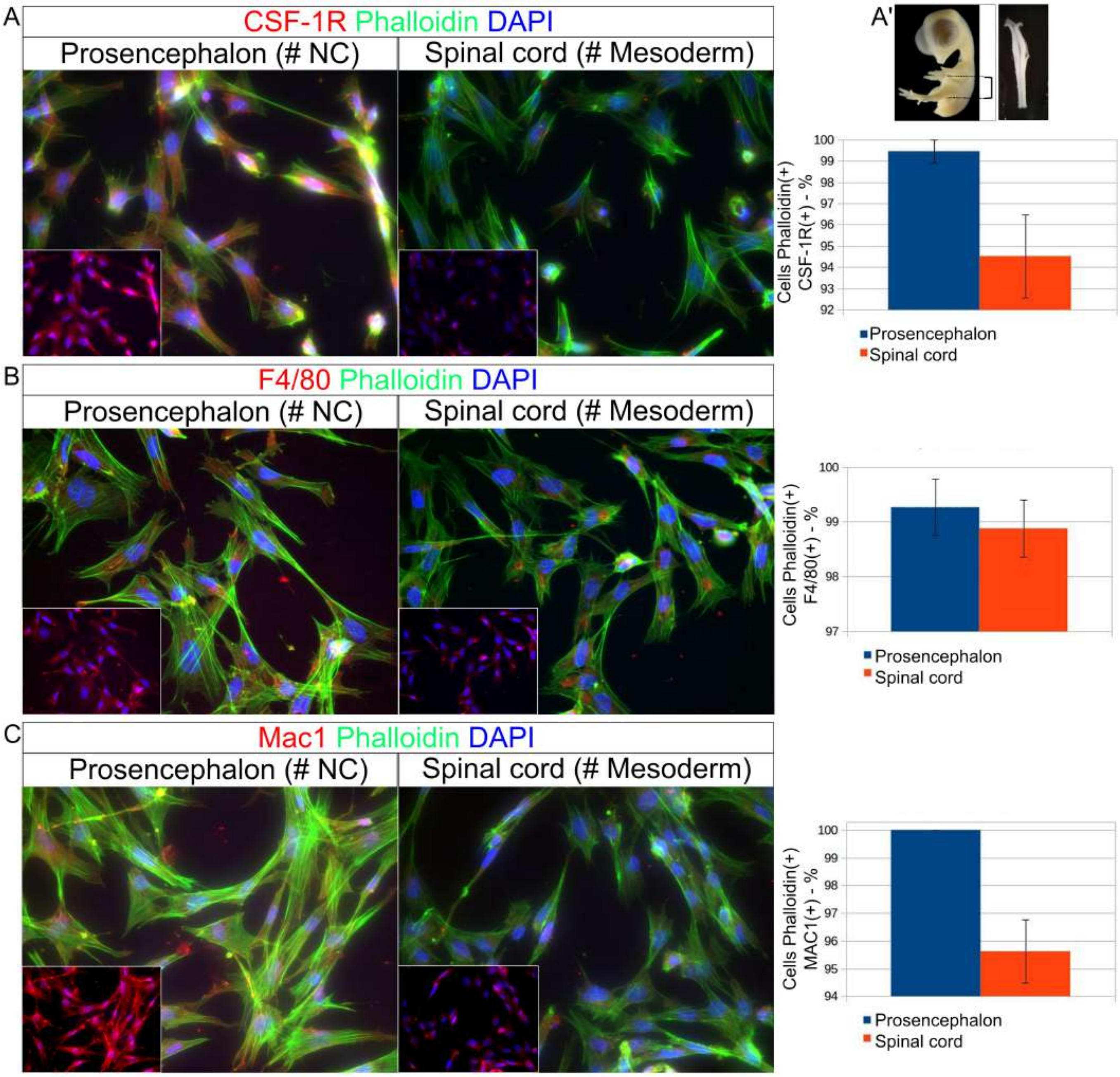

### Supplementary Fig5

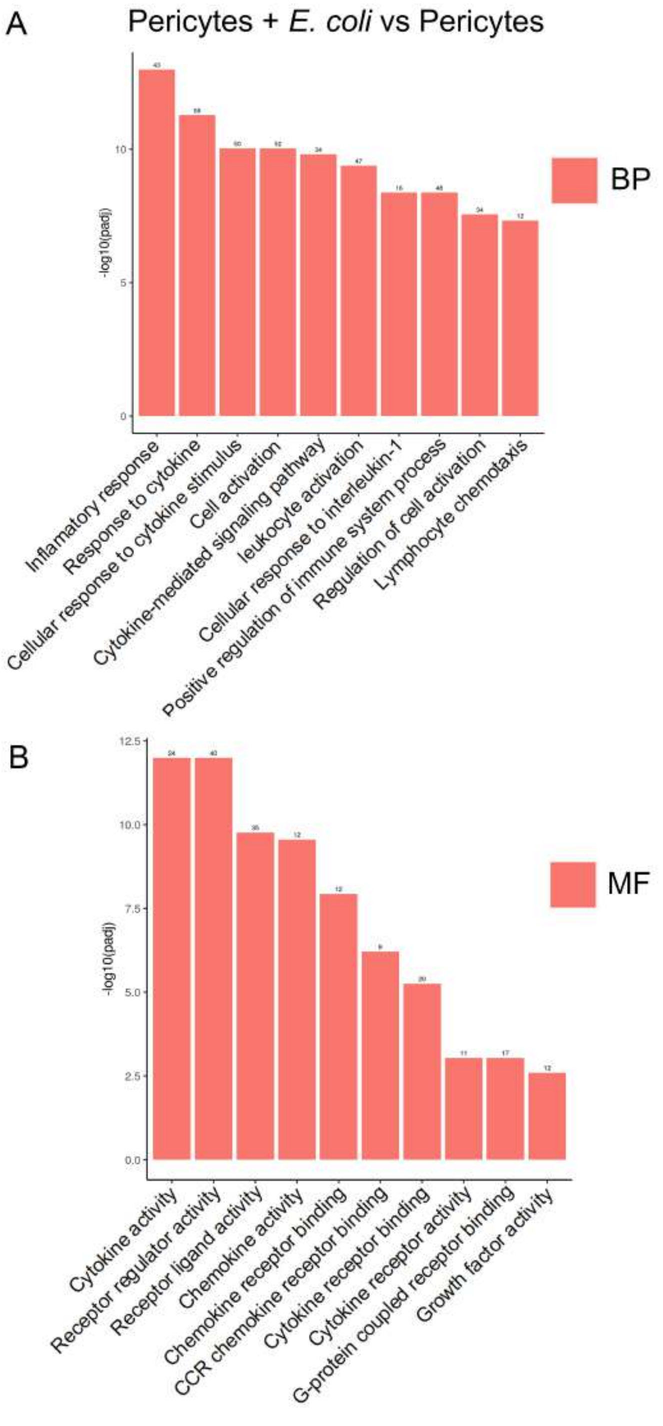

### Supplementary Fig6

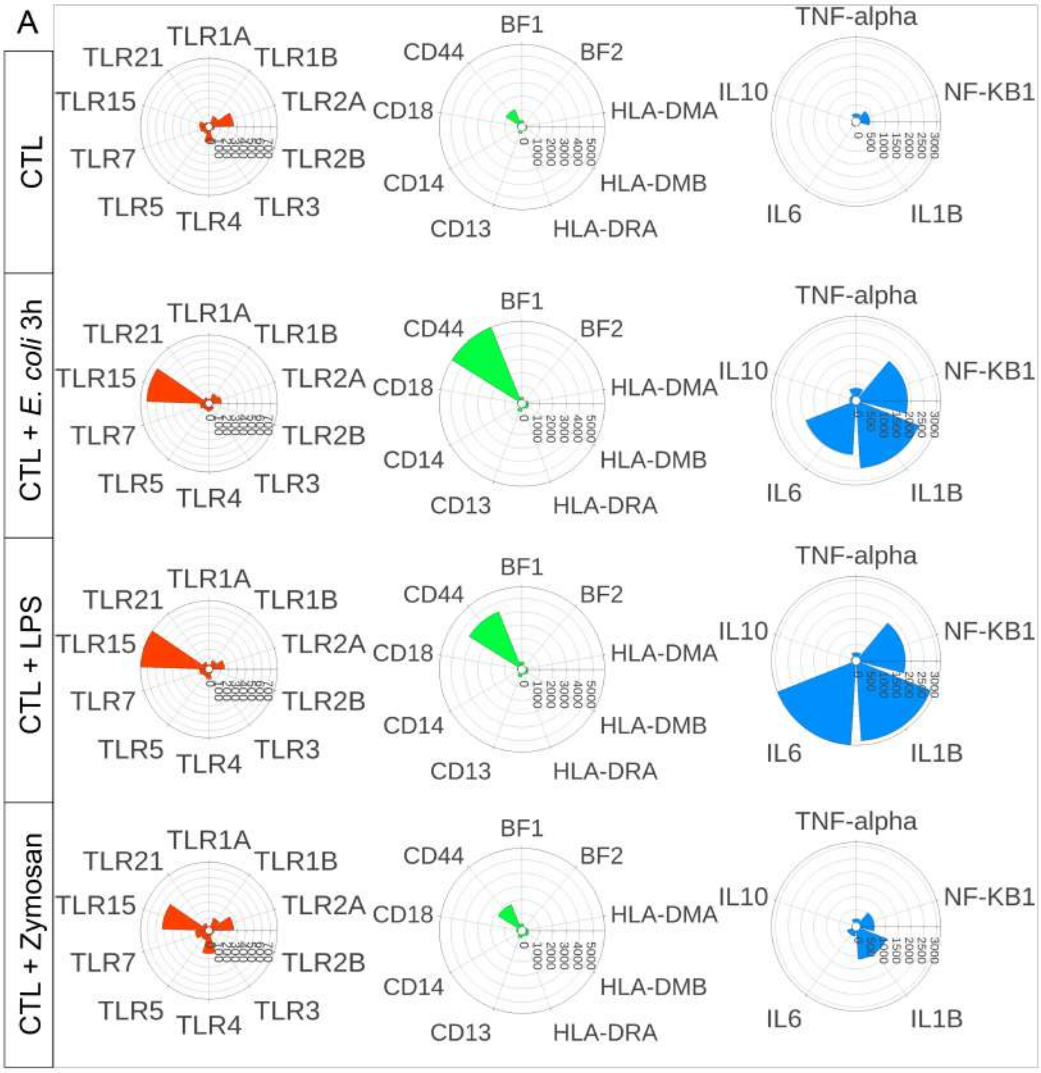

### Supplementary File10

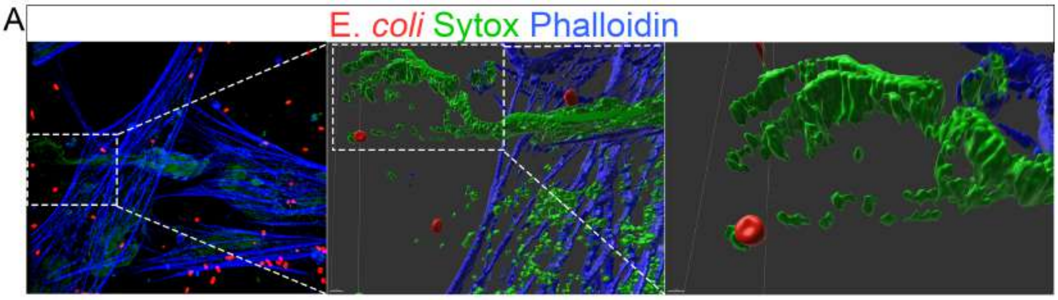
